# A novel mechanism for amplification of sensory responses by the amygdala-TRN projections

**DOI:** 10.1101/623868

**Authors:** Mark Aizenberg, Solymar Rolon Martinez, Tuan Pham, Winnie Rao, Julie Haas, Maria N. Geffen

## Abstract

Many forms of behavior require selective amplification of neuronal representations of relevant environmental signals. Following emotional learning, sensory stimuli drive enhanced responses in the sensory cortex. However, the brain circuits that underlie emotionally driven control of the sensory representations remain poorly understood. Here we identify a novel pathway between the basolateral amygdala (BLA), an emotional learning center in the mouse brain, and the inhibitory nucleus of the thalamus (TRN). We demonstrate that activation of this pathway amplifies sound-evoked activity in the central auditory pathway. Optogenetic activation of BLA suppressed spontaneous, but not tone-evoked activity in the auditory cortex (AC), effectively amplifying tone-evoked responses in AC. Anterograde and retrograde viral tracing identified robust BLA projections terminating at TRN. Optogenetic activation of amygdala-TRN pathway mimicked the effect of direct BLA activation, amplifying tone-evoked responses in the auditory thalamus and cortex. The results are explained by a computational model of the thalamocortical circuitry. In our model, activation of TRN by BLA suppresses spontaneous activity in thalamocortical cells, and as a result, thalamocortical neurons are primed to relay relevant sensory input. These results demonstrate a novel circuit mechanism for shining a neural spotlight on behaviorally relevant signals and provide a potential target for treatment of neuropsychological disorders, in which emotional control of sensory processing is disrupted.

## Introduction

In our everyday experience, we encounter the same sensory stimuli under different behavioral and emotional contexts, which can modify their behavioral relevance. If a stimulus is repeatedly encountered in an emotionally salient context, sensory resources are reallocated to preferentially encode that stimulus *(1, 2)*. This is particularly important for dangerous, fear-evoking stimuli. Behaviorally, the link between emotional learning, such as fear conditioning, and changes in sensory processing, has been established in humans and other mammals. Recently, we found that differential fear conditioning can lead to an impairment or an improvement in sensory discrimination, depending on the generalization of learning, and that the auditory cortex is required for expression of these perceptual changes *(3)*. Similar effects were found following fear conditioning in humans *(4-6)*. Many neuropsychological disorders are characterized by inappropriate emotional weighting of sensory stimuli, including schizophrenia *(7, 8)* and anxiety disorders *(1, 9)*. Untangling the mechanisms that govern emotion-driven control of sensory perception is important not only for basic understanding of sensory processing in everyday environments, but also for identifying potential treatment targets characterized by abnormal emotional responses to benign sensory stimuli.

The baso-lateral amygdala (BLA) is a critically important hub for the formation and expression of fear memories associated with sensory stimuli (for review see *10)*. Aversive stimuli drive strong responses in the BLA *(5, 11, 12) (*although they can be heterogeneous *13)*, and fear conditioning evokes plastic changes in neuronal responses to conditioned sounds in the sensory cortex *(14-17)*. BLA has been proposed to drive plasticity in the auditory cortex for signals associated with fear *(4, 18)*. While changes in the auditory cortex following fear conditioning have been extensively documented (for review see *(17)*, it is not clear whether and how BLA modulates cortical responses to sensory stimuli *(19, 20)*.

Lateral amygdala (LA) sends direct projections to the AC, as evidenced by imaging of LA axons in AC *(21)*. However, recent studies in the primate brain revealed an additional pathway from the BLA to the primary inhibitory nucleus in the thalamus, the thalamic reticular nucleus (TRN) *(22)*. This finding raises the possibility that TRN facilitates gating of signals in the sensory cortex from the BLA. TRN is a layer of inhibitory GABAergic neurons located between neocortex and thalamus, which does not send direct projections to the neocortex but provides inhibition for the sensory thalamocortical relay cells *(23)*. At the same time, TRN receives excitatory collaterals from cortex and thalamus *(24)*. These projections position TRN as a gatekeeper, controlling sensory information flowing from thalamus to cortex, and potentially suppressing irrelevant stimuli, offering an opportunity to reweigh sensory responses based on their behavioral saliency *(25-27)*.

Here, we first tested the effects of activation of BLA on tone-evoked responses in AC. We found that activating BLA suppressed spontaneous activity in AC, leading to an increase in tone-evoked response amplitude. By examining the connectivity between BLA and the thalamus using viral anterograde and retrograde viral tracing techniques in the mouse, we identified direct projections from BLA to TRN. We found that activating this connection selectively suppressed spontaneous activity in the auditory thalamus, similar to the effects of activating BLA on AC. Through a computational model of thalamocortical circuitry, we found that activating BLA inputs to TRN could account for the reduction in spontaneous thalamic activity, and this reduction acted to prime the thalamocortical relay response to sensory input. Together, these findings suggest that the amygdala-TRN pathway amplifies responses to sensory input by suppressing spontaneous activity of relay neurons, a process that could underlie fear-driven changes in auditory and other forms of sensory discrimination.

## Results

### Photo-activation of BLA amplifies tone-evoked responses in the AC

We first tested the effect of optogenetic activation of BLA on spontaneous and tone-evoked activity in AC. To manipulate the level of activity of excitatory neurons in the amygdala, we expressed Channelrhodopsin (ChR2) using targeted viral delivery to BLA of mice expressing Cre recombinase in neurons under the CamKIIα promoter (CamKIIα-Cre mice, Figure 1A). Injection of a modified adeno-associated virus (AAV), which carried the antisense code for ChR2 under the FLEX cassette resulted in efficient and specific expression of ChR2 (flex-ChR2) in excitatory neurons in BLA of CamKIIα-Cre mice (Figure 1B).

**Figure 1.**
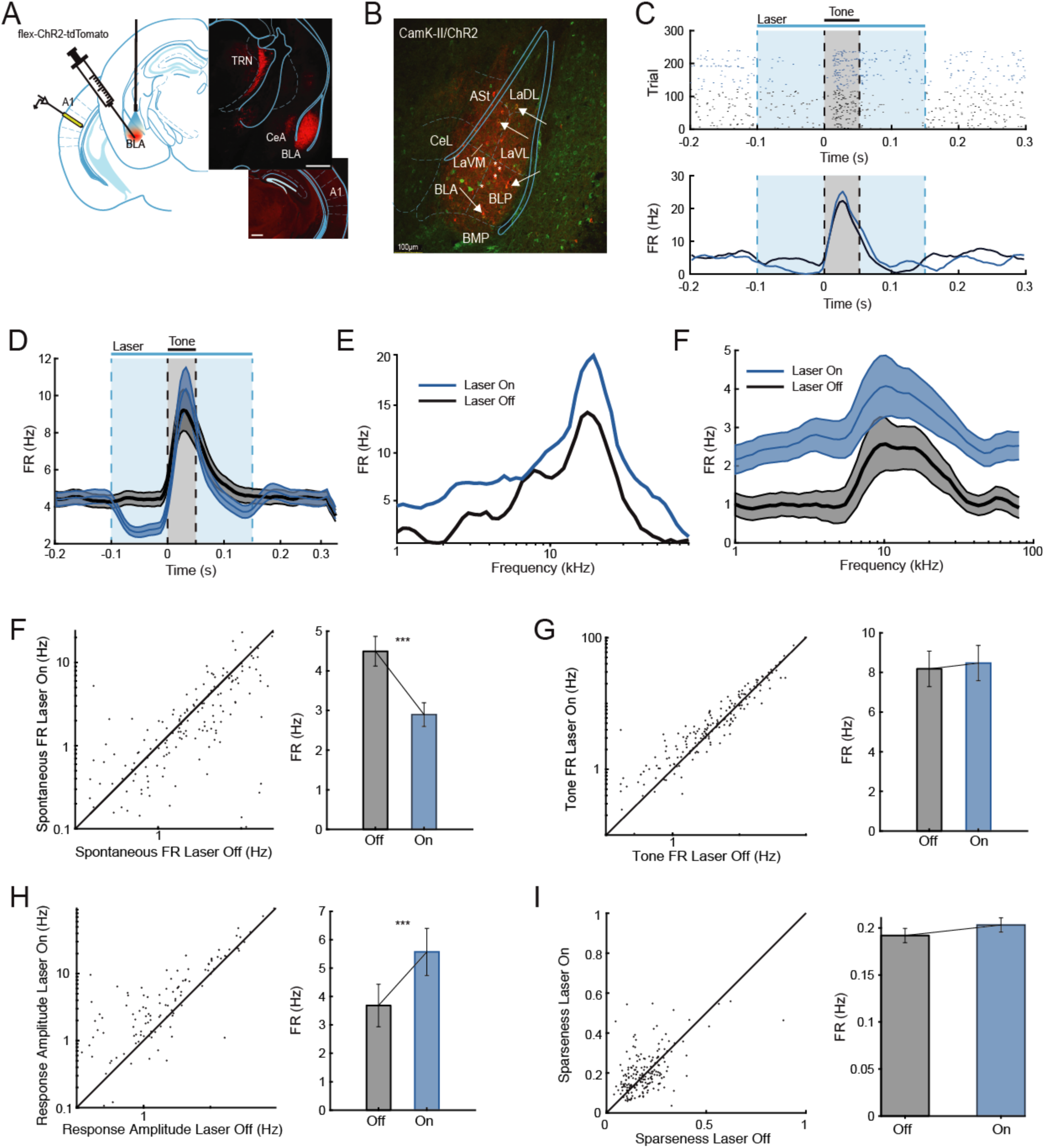
Photo-activation of BLA increases tone-evoked responses in the AC. **A.** Left panel. CamK-cre mice were injected bilaterally with AAV-FLEX-ChR2-tdTomato into BLA. Animals were implanted with optical fibers bilaterally targeting BLA. Putative excitatory neurons in BLA were activated by blue light (473 nm) while neuronal activity in AC was recorded using either a multi-tetrode micro-drive or multichannel silicon probe. Right panel. Top. Micrograph showing expression of injected virus in BLA and its projections to thalamus. Bottom. Micrograph showing little labelling in AC. Scale bar = 0.5 mm. A1: primary auditory cortex; BLA: basolateral amygdala; CeA: central nucleus of amygdala; TRN: thalamic reticular nucleus. **B.** Immunohistochemistry demonstrating co-expression of ChR2-tdTomato in putative excitatory neurons in the BLA of a CamKIIα-Cre mouse. Red: tdTomato. Green: antibody for CamKIIα. Scale bar = 100 microns. **C.** Responses of a representative AC neuron to optogenetic stimulation of BLA. Light was presented from 0 to 0.25 s (blue rectangle). Tone was presented from 0.1 to 0.15 s (grey rectangle). Top: Raster plot of spike times. Bottom. Corresponding peristimulus time histogram (PSTH) of neuronal response in light-On (blue) and light-Off (black) conditions. **D.** PSTH of the AC neurons in response to a tone (grey rectangle) during light-On (blue line) and light-Off (black line) trials. Time of photo-activation of BLA is outlined by a blue rectangle. Plot shows data from all recorded AC neurons (N=190 single units) from 7 mice. Mean ± SEM. **E.** Frequency response function of the neuron from (B) in the absence of photostimulation (Off trials) and during photostimulation of BLA (On trials). **F-I.** Optogenetic activation of BLA suppressed spontaneous firing rate (FRbase, F, paired ttest, t_189_=2.74, p=8.15e-7), but not tone-evoked activity of neurons recorded from AC (FRtone, G, paired ttest, n.s.). Therefore, the amplitude of tone-evoked response was increased (H, paired ttest, t_189_=6.72, p=2.12e-10). Activation of BLA did not affect sparseness of tuning of neurons in AC (I, paired ttest, n.s.). Left panel: Scatter plot of firing rate (F-H) or sparseness (I) on light-On plotted versus light-Off trials. Each circle represents a single unit (red) or multi-unit (black). Right panel: Mean ± SEM of measures from the left panel. ***: p < 0.001 (paired t-test).

We measured spontaneous and tone-evoked activity of neurons in the auditory cortex (AC), targeting the primary auditory cortex (A1), by recording from awake, head-fixed mice during acoustic presentation of a random tone sequence consisting of 50 ms tone pips at 50 frequencies ranging from 1 to 80 kHz (70 dB SPL). BLA was activated by shining blue laser light (473 nm, 3.5 mW/mm^2^ intensity at the fiber tip) through implanted optic cannulas. Photo-activation of BLA significantly reduced overall spontaneous firing rate (FR_base_, computed during the baseline period, 0–50 ms prior to tone onset, N = 190, p = 8.15E-7, df = 189, tstat = 2.74) in AC (Figures 1C, 1D and 1F). While peak tone-evoked firing rate in the AC (FR_tone_, computed 0–50 ms after tone onset) was not significantly affected by BLA activation (Figures 1C, 1D and 1G, n.s.), the amplitude of responses to tones compared with spontaneous firing rate increased (Figures 1D and 1H, N = 190, p = 2.12E-10, tstat= 6.72). Activation of BLA did not significantly affect the tuning bandwidth of neurons in AC, quantified by sparseness of tuning curve (Figures 1E and 1I, n.s.). Thus, by reducing the spontaneous firing rate, but not increasing the tone-evoked response, activation of BLA amplified tone-evoked responses in the auditory cortex.

### BLA sends projections to TRN

Suppression of spontaneous firing in cortical neurons during BLA activation is most likely due to an inhibitory synapse. Recent studies identified direct projections from the BLA to the TRN, an inhibitory nucleus in the thalamus *(22)*. The TRN in turn projects to the MGB, and therefore activation of BLA could drive suppression of spontaneous activity in AC by activating inhibitory TRN to MGB projections. We observed that AAV injected in the BLA resulted in labeling of terminals in the TRN (Figure 1A). To confirm the existence of BLA-TRN pathway, we injected retrograde CAV2-Cre vector in the TRN of AI14 reporter mice, in which tdTomato is conditionally expressed in the cells transfected with Cre recombinase. We identified retrograde labeling of neurons in the basolateral and central nuclei of amygdala (Figure 2A), confirming the existence of a direct pathway from the BLA to the TRN.

**Figure 2.**
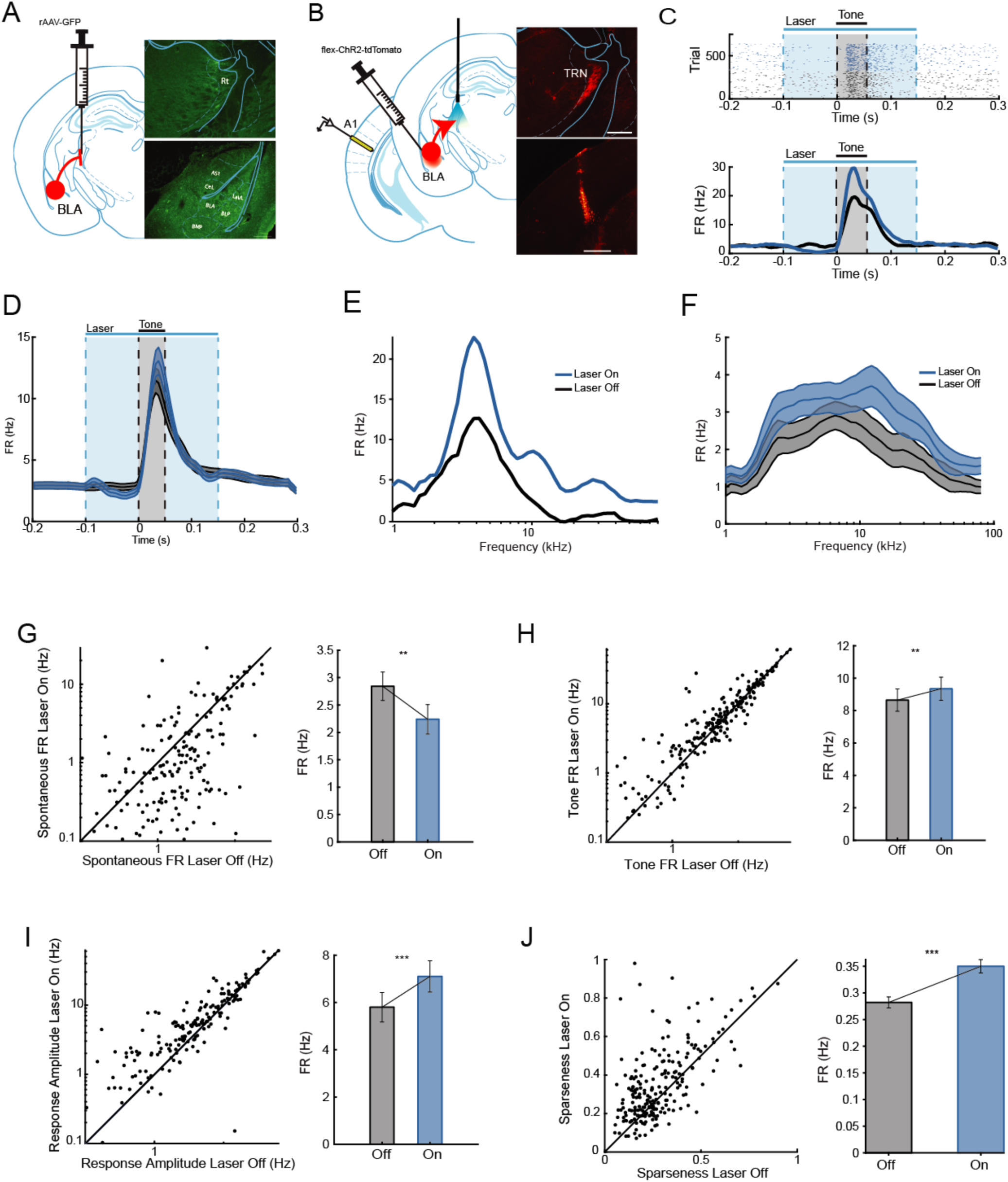
Photo-activation of projections from BLA to TRN increases amplitude of tone-evoked responses in AC. **A.** Left Panel. Retrograde tracing of the BLA projections to TRN. AI14 mice were injected with CAV2-cre in TRN. Right panel. Top: Micrograph showing expression at the viral injection site in TRN. Scale bar = 0.5mm. Bottom: Retrograde expression of tdTomato in neurons in BLA and CeA. Scale bar = 0.1mm. **B.** Anterograde tracing of the BLA projections to TRN. Left panel. Mice were injected bilaterally with the virus expressing ChR2 into BLA. Animals were implanted with optical fibers bilaterally targeting TRN. BLA-TRN projections were activated by blue light (473 nm) on the TRN while neuronal activity in AC was recorded using a multichannel silicon probe. Right panel. Top: Micrograph showing projection terminals in TRN. Bottom: Fluorescent trace from recording silicon probe. Scale bar = 0.5 mm. **C.** Responses of a representative AC neuron to optogenetic stimulation of TRN. Light was presented from 0 to 0.25 s (blue rectangle). Tone was presented from 0.1 to 0.15 s (grey rectangle). Top: Raster plot of spike times. Bottom. Corresponding PSTH of neuronal response in light-On (blue) and light-Off (black) conditions. **D.** Mean PSTH of the AC neurons in response to a tone (grey rectangle) on light-On (blue line) and light-Off (black line) trials. Time of photo-activation of TRN is outlined by a blue rectangle. Plot shows data from all recorded AC neurons (N=216) from 5 mice. Mean ± SEM. **E.** Frequency response function of a representative neuron in the absence of photostimulation (Off trials) and during photostimulation of TRN (On trials). **F-I.** Optogenetic activation of BLA-TRN projections suppressed spontaneous firing rate (FRbase, F, paired ttest, t_215_=2.74, p=0.0067), but increased tone-evoked activity of neurons recorded from AC (G, FRtone, paired ttest, t_215_=2.85, p=0.0048). Amplitude of tone-evoked response was increased (H, paired ttest, t_215_=5.65, p=5.0e-8). Activation of BLA-TRN projections did not affect sparseness of tuning of neurons in AC (I, paired ttest, t_215_=6.62, p=2.73e-10). Left panel: Scatter plot of firing rate (F-H) or sparseness (I) on light-On plotted versus light-Off trials. Each circle represents a single unit. Right panel: Mean ± SEM of measures from the left panel. ***: p < 0.001 (paired t test).

### Photo-activation of BLA-TRN pathway increases amplitude of tone-evoked responses in the AC

To test whether BLA-TRN pathway underlies the amplification of tone-evoked responses caused by BLA activation (Figure 1), we injected either a vector expressing hChR2 under CAG promoter in the BLA of WT mice, or flex-ChR2 vector into the BLA of CamKIIα-Cre mice. We then shined blue laser light onto the TRN through implanted cannulas, while recording neural responses in AC using multichannel silicon probes (Figure 2B). This allowed us to test the effect of selective activation of neurons that project from BLA to TRN on neuronal activity in AC. Activation of BLA terminals in TRN led to a significant suppression of spontaneous neuronal activity in AC (Figure 2C, 2D and 2F, N = 216, p = 0.0067, df = 215, tstat = 2.74), whereas absolute tone-evoked firing rate was significantly increased (Figure 2D and 2G, N = 216, p = 0.0048, df = 215, tstat = 2.85). Hence, average amplitude of responses to tones increased significantly (Figure 2D and 2H, N = 216, p = 5E-8, df = 215, tstat = 5.65). Sparseness of tuning curves decreased as result of TRN activation (Figure 2E and 2I, N = 216, p = 2.73E-10, df = 215, tstat = 6.62).

These effects persisted when we also blocked activity of BLA neurons by focal application of TTX (Supplemental figure S1 1-F, after TTX: N = 74, decrease in spontaneous activity, p = 3.03e-11, df = 73, tstat = 7.8; increase in response amplitude, p = .0061, df = 73, tstat = 2.82; Supplemental figure S1 G, H; before TTX: N = 74, decrease in spontaneous activity, p = 1.49e-10, df = 73, tstat = 7.45; increase in response amplitude, p = 7.01e-6, df = 73, tstat = 4.84). This control ensured that our stimulation of BLA-TRN terminals did not lead to activation of another pathway originating from the cell bodies in the BLA *(28-30)*. Combined, these results demonstrate that selective activation of BLA-TRN projections evokes similar effects to those observed with general BLA activation on activity in the AC.

### Photo-activation of BLA-TRN pathway increases amplitude of tone-evoked responses in the auditory thalamus

There is extensive evidence that TRN inhibits sensory processing in the sensory thalamus *(25, 31)*. Therefore, we hypothesized that effect of BLA-TRN pathway on the AC activity is the result of inhibition that auditory thalamus receives from TRN. We tested whether activation of BLA-TRN terminals drives similar effects to those observed in AC in the auditory thalamus (Medial Geniculate Body, MGB). We tested this hypothesis by optogenetically activating BLA-TRN pathway as described above, while simultaneously recording from MGB (Figure 3A and 3B). Similarly to previous results, photo-activation of amygdalar terminals in TRN led to significant inhibition of spontaneous firing rate of neurons in MGB (Figure 3C and 3E, N = 126, p = 8.97e-10, df = 189, tstat = 5.1). In contrast, mean tone-evoked activity was not affected by photo-stimulation (Figure 3C and 3F, n.s.), resulting in increased amplitude of tone-evoked responses (Figure 3G, p = 9.6e-7, df = 125, tstat = 5.16). Similarly to AC recordings, photo-activation of BLA-TRN pathway increased the sparseness of tuning curve (Figure 3D and 3H, p = 8.65 e-5, df = 125, tstat = 4.06). Combined, our findings are consistent with the hypothesis that BLA-TRN pathway acts on the AC through MGB.

**Figure 3.**
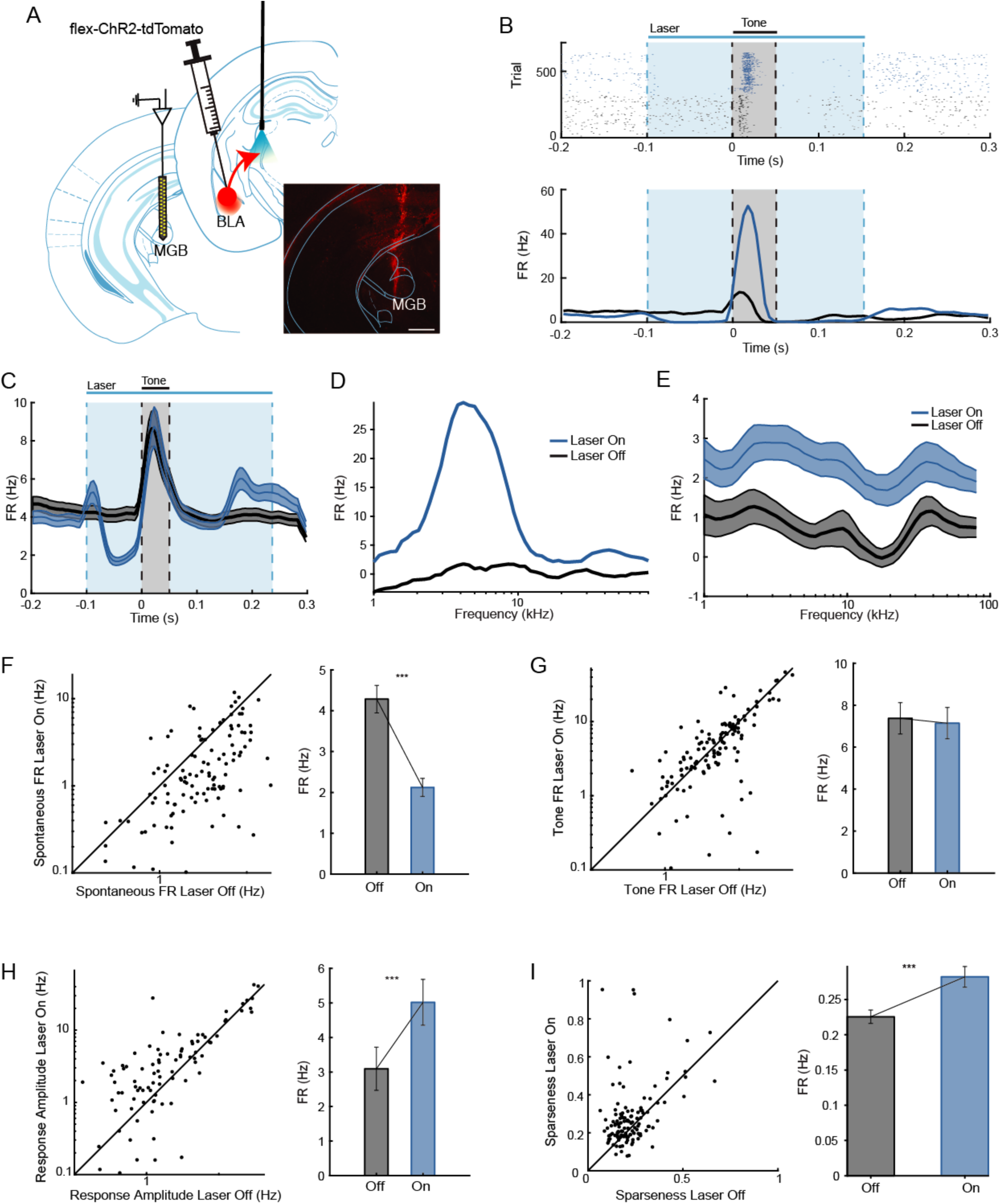
Photo-activation of projections from BLA to TRN increases amplitude of tone-evoked responses in the MGB. **A.** Left panel. Mice were injected bilaterally with a virus expressing ChR2 into BLA. Animals were implanted with optical fibers bilaterally targeting TRN. BLA-TRN projections were activated by blue light (473 nm) while neuronal activity in MGB was recorded using a multichannel silicon probe. Right panel. Micrograph showing fluorescent trace from the recording silicon probe. Scale bar = 0.5 mm. **B.** Responses of a representative MGB neuron to optogenetic stimulation of TRN. Light was presented from 0 to 0.25 s (blue rectangle). Tone was presented from 0.1 to 0.15 s (grey rectangle). Top: Raster plot of spike times. Bottom. Corresponding PSTH of neuronal response in light-On (blue) and light-Off (black) conditions. **C.** PSTH of the MGB neurons in response to a tone (grey rectangle) during light-On (blue line) and light-Off (black line) trials. Time of photo-activation of TRN is outlined by a blue rectangle. Plot shows data from all recorded MGB neurons (N=126) from 5 mice. Mean ± SEM. **D.** Frequency response function of the neuron from (B) in the absence of photostimulation (Off trials) and during photostimulation of TRN (On trials). **E-H.** Optogenetic activation of BLA-TRN projections suppressed spontaneous firing rate (FRbase, E, paired ttest, t_125_=5.1, p=8.97e-10), but did not significantly change tone-evoked activity of neurons recorded from MGB (FRtone, F, paired ttest, n.s.). The amplitude of tone-evoked response was increased (G, paired ttest, t_125_=5.16, p=9.6e-7. Activation of BLA-TRN projections increased the sparseness of tuning of neurons in MGB (H, paired ttest, t_125_=4.06, p=8.65e-5). Left panel: Scatter plot of firing rate (E-G) or sparseness (H) on light-On plotted versus light-Off trials. Each circle represents a single unit. Right panel: Mean ± SEM of measures from the left panel. ***: p < 0.001 (paired t test).

To verify that effect of light on the activity in auditory thalamus and cortex is specific to action of ChR2, we injected control group of mice with viral vector that encoded only fluorescent protein. In control mice, shining blue light on BLA projections in TRN did not cause any significant change in the firing rates of neuron either in AC (Figure S2) or in MGB (Figure S3).

### A bursting model of TRN and MGB produces enhancement in tone-evoked responses in TRN and A1 due to BLA-TRN activation

How does the enhanced responsiveness to tones square with the decreased spontaneous activity upon BLA-TRN activation? These dual effects may be explained by the feedback connectivity between the bursting neurons in TRN and MGB. We constructed a three-cell model of MGB, TRN and A1 neurons. As all thalamic neurons exhibit bursts of spikes resulting from T currents *(32-34)*, we used a bursting neuron model for our TRN and MGB neurons *(35, 36)*. Spontaneous spiking was elicited by Poisson-distributed inputs to the MGB and A1 neurons. We delivered a 100-ms long pulse of input to the TRN neuron to represent optogenetic activation of BLA terminals in TRN, and a shorter 10-ms input to MGB to represent the tone input. BLA activation initiated a burst in TRN neurons, with regular tonic firing following the burst for increased strengths of BLA activation (Fig. 4B1). For weak BLA input, the initial burst in TRN was late and elicited fewer spikes, providing only weak inhibition of the spontaneous firing in MGB and of the MGB response when the tone arrived (second row in Fig. 4 B2-3 and C). For moderate values of BLA activation, a quicker and stronger burst in TRN (Fig. 4 C1) delivered stronger and earlier inhibition in MGB and A1 (third row in Fig. 4B2-3 and C2-3). This inhibition terminated spontaneous firing in MGB, and the delayed timing of the TRN burst resulted in stronger rebound from inhibition in the MGB neuron. Together, these effects allowed MGB and A1 to produce an amplified response when the tone arrived. Overactivation of BLA (bottom row in Fig. 4B2-3 and C2-3) led to overactivation of TRN and correspondingly stronger and persistent inhibition of MGB and A1. These effects of strength and timing of TRN bursts on firing rates in MGB and A1 are summarized in Fig. 4D. Consistent with the experimental results, the suppression of spontaneous MGB spiking allowed for an increased ability to respond to the auditory input and relay it to A1. Together, these results provide a link between the suppression of spontaneous activity to amplified A1 response by demonstrating that a moderate activation by BLA of TRN, such as those used in our experiments, increase the readiness of MGB neurons to respond. As a result, peak-to-peak firing rates in MGB and AC increased during BLA activation (Fig. 4E).

**Figure 4.**
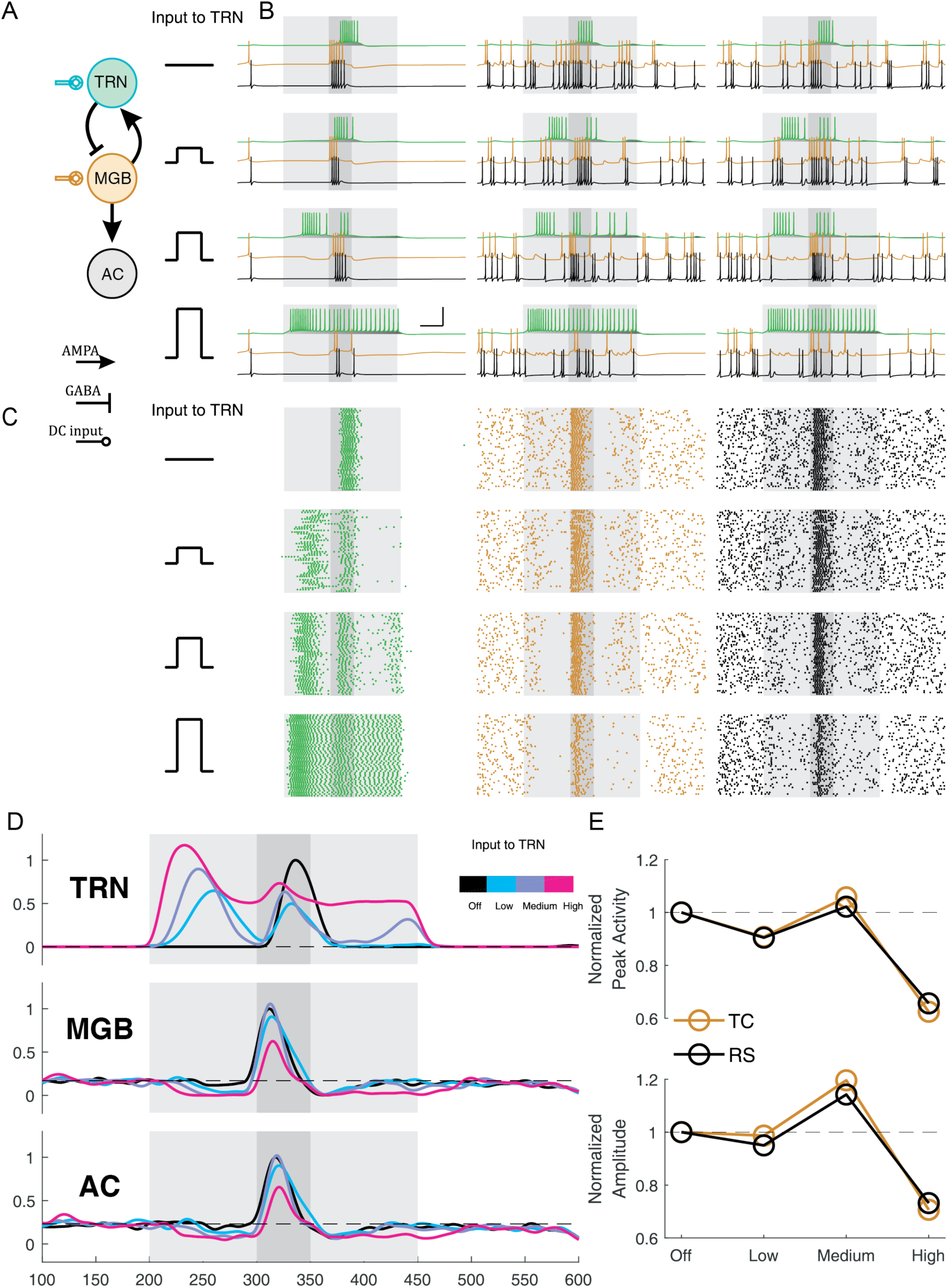
Possible mechanism for BLA enhancement of tone-evoked MGB and A1 responses. **A.** Simulated network, including TRN, MGB and A1 cells (see Methods). A 100-ms input to TRN represented optogenetic activation of BLA. The tone was a 10-ms pulse to MGB during light stimulus. **B.** Simulations of the network for no spontaneous inputs (B_1_) and for two simulations with spontaneous input to MGB and A1 (B_2-3_), in the absence of BLA activation (top row) and with increasing strengths of BLA activation (lower rows). **C.** Spike rasters for 25 repeated simulations in **B. D**. Spiking rates in the TRN, MGB and A1 neurons, shown for control conditions (black) and for the three strengths of BLA activation of TRN (shades of purple), each normalized to zero-input values. **E.** Maximum firing rate in MGB and A1 during the tone (E_1_), and peak-to-peak amplitude of the tone response (E_2_) shown for varied strength of BLA activation.

## Discussion

The basolateral amygdala is a critically important brain area for auditory fear conditioning (for review see *(10)*, where conditioned and unconditioned stimuli converge *(37, 38)*. Lesions of the BLA *(39,40)* or either the lemniscal or non-lemniscal auditory inputs to the BLA, impair acquisition and expression and discrimination of conditioned fear responses *(41, 42)*; whereas paired activation of the BLA with an auditory cue is sufficient to induce a conditioned fear response *(43)*. Recent studies found that sensory fear conditioning can modulate sensory discrimination acuity *(4, 6)*, and we demonstrated that in the auditory system this modulation requires the auditory cortex *(44)*. Therefore, in the present study, we examined whether and how activating BLA affects tone-evoked responses in AC. We first found that activating BLA increases tone-evoked response amplitude in AC by suppressing spontaneous activity, but not affecting tone-evoked responses. We identified a mechanism by which this suppression can occur: via the projections from the BLA to the inhibitory nucleus of the thalamus, the TRN. Consistent with established anatomy, we did not find significant direct projections from TRN to AC. Therefore, we hypothesized that the TRN controls responses in AC via the MGB of the auditory thalamus *(45)*. Indeed, specifically activating BLA-TRN projection neurons drove an increase in tone-evoked response amplitude in both MGB and AC. The effects were stronger in MGB than AC further suggesting that the MGB serves as a relay for cortico-collicular control. These effects could be accounted for by a three-cell model of the TRN-MGB-AC connections (Figure 4), with the critical effect provided by the difference in timing and magnitude of the inhibition that TRN delivered to MGB, consistent with previous thalamo-cortical models *(33, 34, 36, 46)*. This study thus establishes an important pathway connecting the emotional and sensory processing centers that potentially drives changes in auditory perception as result of emotional learning.

The thalamic reticular nucleus is a thin sheet of GABAergic neurons surrounding dorsal thalamus, which exhibits sectorial anatomical organization, such that each sector of the TRN is specific to a sensory modality *(47, 48)*. Although the TRN does not send direct projections to sensory cortical areas, it can control the flow of auditory and other sensory information to the cortex by inhibiting or disinhibiting thalamic projection neurons in the auditory thalamus *(45, 49-51)*. The unique anatomical and functional organization of TRN gave rise to the “attentional searchlight” hypothesis *(52)*, which proposed that the TRN drives attention toward salient stimuli, by inhibiting sensory responses to irrelevant information. Our results imply that that BLA is one of the controls of that searchlight, control that is exerted by inhibiting spontaneous activity in the relay cells. Our parameters also suggest that fear-driven BLA activation that is too weak, or conversely overwhelming, fail to control the spotlight. We suggest that this simple mechanism may apply to multiple arising senses as they pass through the thalamocortical circuit.

Communication across the TRN *(53)* or by convergence and divergence of TRN-thalamic connections offers the possibility for the activation of one specific sense, or sensory modality, to affect the thalamic relay of another sense *(33, 54)*. Recently, the TRN was experimentally shown to selectively amplify processing of task-relevant stimuli and attention-guided behaviors. Either genetic (knockout or knockdown of the ErbB4 receptor in TRN neurons) or optogenetic perturbation of neuronal activity in the TRN diminished attentional switching between conflicting sensory cues in a two-alternative choice task *(25, 27)*. Similarly, optogenetic activation of the TRN during the window of elevated attention to a visual cue interfered with performance in a visual detection task *(26)*. Our present results showing that TRN activity is modulated by its inputs from BLA suggest that emotional responses generated in the amygdala may also modulate sensory interactions within and through the TRN, particularly during fear learning.

Auditory fear conditioning drives plastic changes to tone-evoked responses in auditory thalamus *(55)* and auditory cortex *(17, 56)*. Multi-neuronal recording in AC demonstrated that tone-evoked responses to the conditioned stimulus are increased following fear conditioning *(57)*, with individual neurons exhibiting heterogeneous but sustained changes in their tuning properties *(56)*. Auditory fear conditioning promotes formation of dendritic spines in AC *(58)*, pointing to plastic changes in neuronal connectivity. Direct amygdala-cortical projections are thought to underlie the facilitation of responses to emotionally salient stimuli *(21, 59, 60)*, as fear conditioning leads to an increase in the post-synaptic spines and pre-synaptic boutons specific to BLA-AC neuronal pairs *(21)*. Here, we demonstrate that a parallel processing pathway to cortex from the BLA, via the TRN and MGB (Figure 5). This pathway may potentially play a regulatory role during the acquisition and recall of auditory fear memories. These two pathways may complement each other in enhancing responses to the conditioned stimulus by strengthening amygdala-cortical connectivity. In future studies, it will be important account for the interactions between these pathways in interpreting the effects of fear conditioning and learning on sensory responses in the cortex.

**Figure 5.**
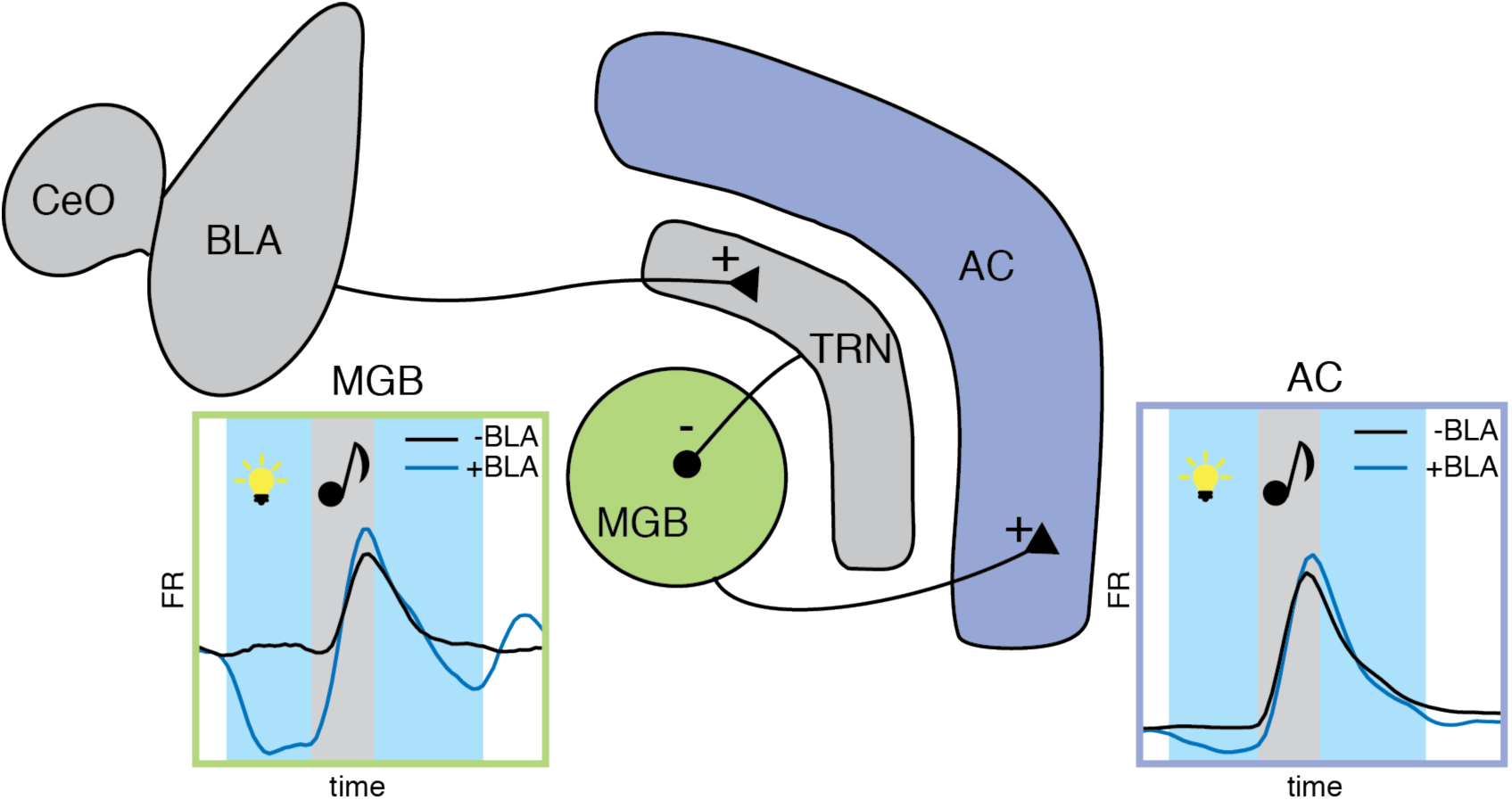
Diagram showing proposed circuit underlying the effects of the amygdala-TRN pathway on auditory processing. Photo-activation of BLA-TRN projections (‘+’ synapses onto the TRN) leads to inhibition of spontaneous activity and amplification of the amplitude of tone-evoked activity of MGB neurons as result of inhibition from TRN (‘-’ synapses onto MGB). This, in turn, amplifies auditory responses in the auditory cortex (‘+’ synapses onto AC). Inserts: representative tone-evoked responses in MGB and AC. Blue lines: during BLA-TRN activation. Black lines: baseline. Gray: tone; blue: laser on.

We previously found that generalized auditory fear learning led to an impairment in frequency discrimination acuity, whereas specialized learning led to an improvement in acuity *(44)*. Similar bi-directional changes in auditory discrimination were achieved by manipulating the activity of inhibitory interneurons in the cortex *(3)*. The existence of parallel pathways for controlling tone-evoked responses after activation of BLA may be useful for enabling bi-directional changes in sensory processing. In particular, top-down control of inhibition early in sensory processing is particularly useful in gating incoming sensory information. The connection that we identified here may be a manifestation of a more general principle of control of behavioral performance via inhibitory-excitatory interactions *(61)*.

## Methods

### Animals

All experiments were performed in adult female (N=5) or male (N=14) mice (supplier: Jackson Laboratories) between 7–15 weeks of age weighing between 17–27 grams. Strains: CamKIIα-Cre: B6. Cg-Tg(CamKIIα-Cre)T29-1Stl/J; wild-type mice: C57BL/6J), AI14: Rosa-CAG-LSL-tdTomato-WPRE::deltaNeo housed at 28°C on a 12 h light – dark cycle with water and food provided ad libitum, with fewer than five animals per cage. In CamKIIα-Cre mice, Cre was expressed in excitatory neurons. All animal work was conducted according to the guidelines of University of Pennsylvania IACUC and the AALAC Guide on Animal Research. Anesthesia by iso-fluorane and euthanasia by carbon dioxide were used. All means were taken to minimize the pain or discomfort of the animals during and following the experiments.

### Surgery and Virus Injection

At least 10 days prior to the start of experiments, mice were anesthetized with isoflurane to a surgical plane. The head was secured in a stereotactic holder. For recordings targeting AC, the mouse was subjected to a small craniotomy (2 × 2 mm) over left AC under aseptic conditions (coordinates relative to Bregma: −2.6 mm anterior, 4.2 mm lateral, +1 mm ventral). For recordings targeting MGN, the mouse was subjected to a small craniotomy (0.5 x 0.5 mm) over left MGN (coordinates relative to Bregma: −3.2 mm anterior, 2.0 mm lateral). For optogenetic activation of BLA neurons, a small craniotomy (0.5 × 0.5 mm) was performed bilaterally over amygdala (coordinates relative to Bregma: 1.5 mm posterior, ±3.0 mm lateral). Fiber-optic cannulas (Thorlabs, Ø200 μm Core, 0.39 NA) were implanted bilaterally over the craniotomy at depth of 4.4 mm from the Bregma. For optogenetic activation of BLA projections to TRN, a small craniotomy (0.5 × 0.5 mm) was performed bilaterally over TRN (coordinates relative to Bregma: −1.1 mm anterior, ±2.0 mm lateral). Fiber-optic cannulas were implanted bilaterally at depth of 2.7 mm from Bregma. Viral constructs were injected using a syringe pump (Pump 11 Elite, Harvard Apparatus) either in BLA (200-400 nl, 4.6 mm depth from Bregma) or in TRN (200 nl, 3.4 mm depth from Bregma). Craniotomies were covered with a removable silicon plug. A small head-post was secured to the skull with dental cement (C&B Metabond) and acrylic (Lang Dental).

For postoperative analgesia, Buprenex (0.1 mg/kg) was injected intraperitonially and lidocaine was applied topically to the surgical site. An antibiotic (0.3% Gentamicin sulfate) was applied daily (for 4 days) to the surgical site during recovery. Virus spread was confirmed postmortem by visualization of fluorescent protein expression in fixed brain slices, and its co-localization with excitatory neurons, following immuno-histochemical processing with the anti-CAMKIIα antibody.

### Viral Vectors

Modified AAV vectors were obtained from Penn VectorCore. Modified AAV encoding ChR2 under FLEX promoter (Addgene plasmid 18917 AAV-FLEX-ChR2-tdTomato) was used for activation of excitatory neurons in CamKIIα-Cre mice. hChR2 was used for activation of neurons in WT mice (Addgene 20938M AAV5-CAG-hChR2(H134R)-mCherry-WPRE-SV40). Modified AAV vectors encoding only tdTomato under a FLEX cassette were used as a control for the specific action of ChR2 on the neuronal populations. Cav2-cre virus (Viral Vector Production Unit) was used for retrograde tracing of BLA-TRN projections in AI24 mice that express tdTomato under FLEX cassette.

### Histology

Brains were extracted following perfusion of 0.01 M phosphate buffer pH 7.4 (PBS) and 4% paraformaldehyde (PFA), postfixed in PFA overnight and cryoprotected in 30% sucrose. Free-floating coronal sections (40 μm) were cut using a cryostat (Leica CM1860). Sections were washed in PBS containing 0.1% Triton X-100 (PBST; 3 washes, 5 min), incubated at room temperature in blocking solution (10% normal horse serum and 0.2% bovine serum albumin in PBST; 1h), and then incubated in primary antibody diluted in carrier solution (1% normal horse serum and 0.2% bovine serum albumin in PBST) overnight at 4°C. Anti-CAMKIIα antibody was used to stain excitatory neurons (abcam5683 rabbit polyclonal, 1:500, abcam). The following day sections were washed in PBST (3 washes, 5 min), incubated for 2 hours at room temperature with secondary antibodies (Alexa 488 goat anti-rabbit IgG; 1:750), and then washed in PBST (4 washes, 10 min). Sections were mounted using fluoromount-G (Southern Biotech) and confocal or fluorescent images were acquired (Leica SP5 or Olympus BX43)

### Photostimulation of Neuronal Activity

Neurons were stimulated by application of five 25 ms-long light pulses (25 ms inter-pulse interval) of blue laser light (473 nm, BL473T3-150, used for ChR2 stimulation), delivered through implanted cannulas. Timing of the light pulse was controlled with microsecond precision via a custom control shutter system, synchronized to the acoustic stimulus delivery. Prior to the start of the experiment, the intensity of the blue laser was adjusted to 3.5 mW/mm^2^ as measured at the tip of the optic fiber.

### Electrophysiological Recordings

All recordings were carried out as previously described *(3)* inside a double-walled acoustic isolation booth (Industrial Acoustics). Mice were placed in the recording chamber, and their headpost was secured to a custom base, immobilizing the head. Activity of neurons was recorded either via a 32-channel silicon multi-channel probe (Neuronexus), or custom-made Microdrive housing multiple tetrodes lowered into the targeted area via a stereotactic instrument following a durotomy. Electro-physiological data were filtered between 600 and 6000 Hz (spike responses), digitized at 32kHz and stored for offline analysis (Neuralynx). Spikes belonging to single neurons were sorted using commercial software (Plexon).

### Acoustic Stimulus

Stimulus was delivered via a magnetic speaker (Tucker-David Technologies), calibrated with a Bruel and Kjaer microphone at the point of the subject’s ear, at frequencies between 1 and 80 kHz to ± 3 dB. To measure the frequency tuning curves, we presented a train of 50 pure tones of frequencies spaced logarithmically between 1 and 80 kHz, at 70 dB, each tone repeated twice in pseudo-random sequence, counter-balanced for laser presentation. The full stimulus was repeated 5 times. Each tone was 50 ms long, with inter-stimulus interval (ISI) of 450 ms. Laser stimulation occurred during every other tone, with an onset 100 ms prior to tone onset. Laser stimulation on each trial consisted of five 25 ms-long pulses with 25 ms-long inter-pulse intervals.

### Neuronal Response Analysis

The spontaneous firing rate (FRbase) was computed from the average firing rate 50 ms before tone onset for light-On and light-Off trials. The tone-evoked firing rate (FRtone) was computed as the average firing rate from 0 to 50ms after tone onset. To examine frequency selectivity of neurons, sparseness of frequency tuning was computed as:

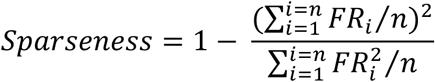

where FR_i_ is tone-evoked response to tone at frequency i, and n is number of frequencies used.

The amplitude of neuronal response to tones was defined as the difference between mean spontaneous (0–50 ms before tone onset) and tone-evoked (0–50 ms after tone onset) firing rate and, for each neuron. Only responses to tones within 0.5 octaves of best frequency (the frequency which resulted in maximum firing rate) of each neuron were included.

### Statistical Analysis

Data were analyzed using two-tailed paired t-tests in Matlab (Mathworks).

### Modeling

We used single-compartment Hodgkin-Huxley neuron models to create a 3-cell network consisting of a thalamic reticular nucleus (TRN) neuron, a thalamocortical neuron representing the MGB and a regular spiking neuron representing primary auditory cortex (A1; Fig. 4A). Starting from the mechanistic models of Traub et al. (2005) and Haas and Landisman (2012), we tuned cells with the following characteristics (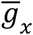 are maximal conductances with units of [*mS*/*cm*^2^] and *E*_*x*_ are reversal potentials in [*mV*]). We used the NEURON implementation in ModelDB of Traub et al. (2005).

| | $\bar{g}_L$ | $\bar{g}_{Naf}$ | $\bar{g}_{Kdr}$ | $\bar{g}_{KA}$ | $\bar{g}_{K2}$ | $\bar{g}_{CaT}$ | $\bar{g}_{AR}$ | $E_K$ | $E_L$ | $E_{Na}$ | $E_{Ca}$ | $E_{AR}$ |
| --- | --- | --- | --- | --- | --- | --- | --- | --- | --- | --- | --- | --- |
| TRN | 0.1 | 60.5 | 60 | 5 | 0.5 | 0.75 | 0.025 | -100 | -75 | 50 | 125 | -40 |
| TC | 0.04 | 100 | 33.75 | 6 | 2 | 0.75 | 0.25 | -95 | -70 | 50 | 125 | -35 |
| RS | 0.04 | 200 | 170 | 20 | 0.5 | 0.1 | 0.1 | -95 | -70 | 50 | 125 | -35 |

Chemical synapses included fast inhibitory GABA_A_ (*E*_*GABAR*_ = –80 *mV*) and excitatory AMPA (*E*_*AMPAR*_ = 0 *mV*) synapses, both with NEURON implementation of AMPA point process synapses, in which the postsynaptic potentials consist of both rise time and fall times, with the former being 0.999 of the latter (ModelDB***). In our simulated network, MGB sent a feedforward excitatory AMPA synapse to TRN with 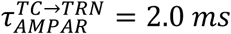 and 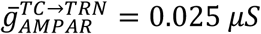. The TRN neuron sent feedback inhibition via a GABA_A_ synapse to the MGB neuron, with both fast and slow fall times (3.3ms and 10ms respectively *(35)*) each contributing equally to the GABA_A_ conductance 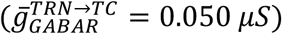. Finally, MGB also sent an AMPA synapse to A1 with 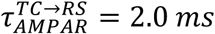 and 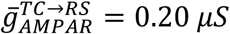. We set the synaptic delay to be 2.0 *ms* and the event detection threshold to be 25 *mV*.

We simulated the network in NEURON for 600 *ms*, with *dt* = 0.005*ms, V*_0_ = –60*mV* and saved sampled data for visualization (Fig 4B) with sampling rate of 0.1*ms*. To simulate spontaneous activity in MGB, we added AMPA synapses with Poisson inputs, where 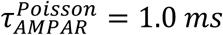 and 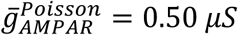, to MGB at 50 Hz, to A1 at 20 Hz and to TRN at 1 Hz. We used holding current to MGB at 1nA and to A1 at 0.5nA. We ran one simulation without any Poisson inputs (Fig. 4B_1_) and 50 simulations for each condition with random Poisson inputs (Fig 4B_2-3_, Fig 4C). In all simulations, we delivered a DC input of 10nA to MGB representing the tone inputs. We delivered a 100-ms input to TRN to represent BLA activation, at three strengths (0 nA, 0.5 nA, 0.8 nA, and 1.8 nA).

To quantify the results of simulations, we calculated the histograms of spike times, binned at 1 ms, then smoothed with a Hanning window of size 31. We normalized each rate to the maximum rate in the control condition preceding input to TRN. To calculate peak activity, we obtained the raw peak activity in the windows of tone input to MGB input for MGB and A1, then normalized those values to the control conditions. Peak-peak amplitude was taken as the difference between the raw peak activity during the tone and the mean activity during the 50 ms before the tone, also normalized to the control condition.

**Figure S1.**
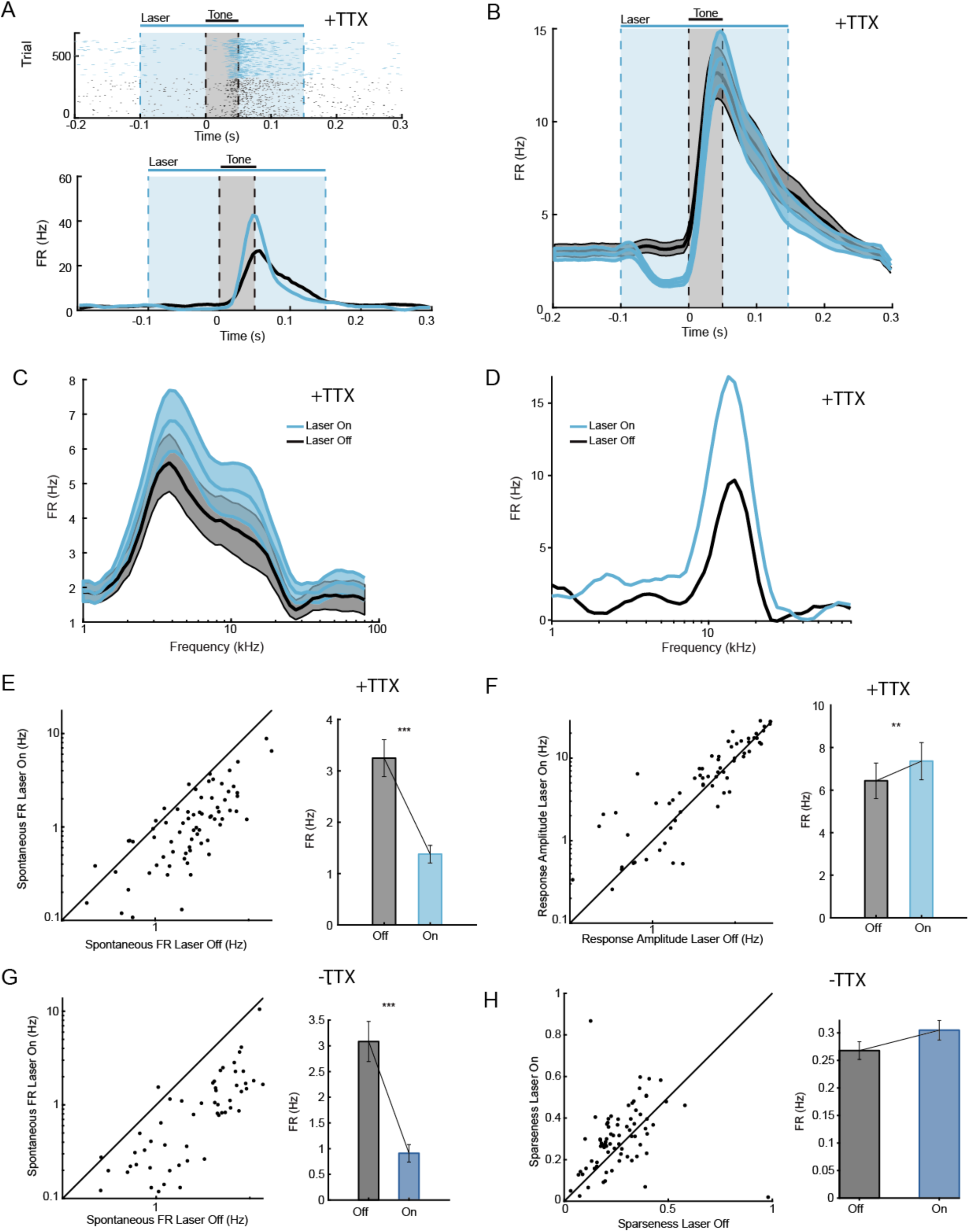
(Supplemental to Figure 2). Photo-activation of projections from BLA to TRN concurrent with administration of TTX in BLA increases amplitude of tone-evoked responses in AC. **A.** Responses of a representative AC neuron to optogenetic stimulation of TRN during concurrent administration of TTX. Light was presented from 0 to 0.25 s (blue rectangle). Tone was presented from 0.1 to 0.15 s (grey rectangle). Top: Raster plot of spike times. Bottom. Corresponding PSTH of neuronal response in light-On (blue) and light-Off (black) conditions. **B.** Mean PSTH of the AC neurons in response to a tone (grey rectangle) on light-On (blue line) and light-Off (black line) trials during concurrent administration of TTX. Time of photo-activation of TRN is outlined by a blue rectangle. Plot shows data from all recorded AC neurons (N = 74) from 2 mice. Mean ± SEM. **C.** Frequency response function from all recorded neurons during concurrent administration of TTX. **D.** Frequency response function of a representative neuron in the absence of photostimulation (Off trials) and during photostimulation of TRN (On trials). **E, F.** Optogenetic activation of BLA-TRN projections during concurrent administration of TTX suppressed spontaneous firing rate (FRbase, F, paired ttest, t_73_=7.9, p=3.03e-11), but increased tone-evoked activity of neurons recorded from AC (G, FRtone, paired ttest, t_73_=2.82, p=0.0061). **G, H.** Optogenetic activation BLA-TRN projections in these mice prior to administration of TTX had the same effects: suppressed spontaneous firing rate (FRbase, F, paired ttest, t_73_=7.45, p=1.49e-10), but increased tone-evoked activity of neurons recorded from AC (G, FRtone, paired ttest, t_73_=4.84, p=7.01e-6). Left panel: Scatter plot of firing rate (E-H) on light-On plotted versus light-Off trials. Each circle represents a single unit. Right panel: Mean ± SEM of measures from the left panel. ***: p < 0.001 (paired t test).

**Figure S2.**
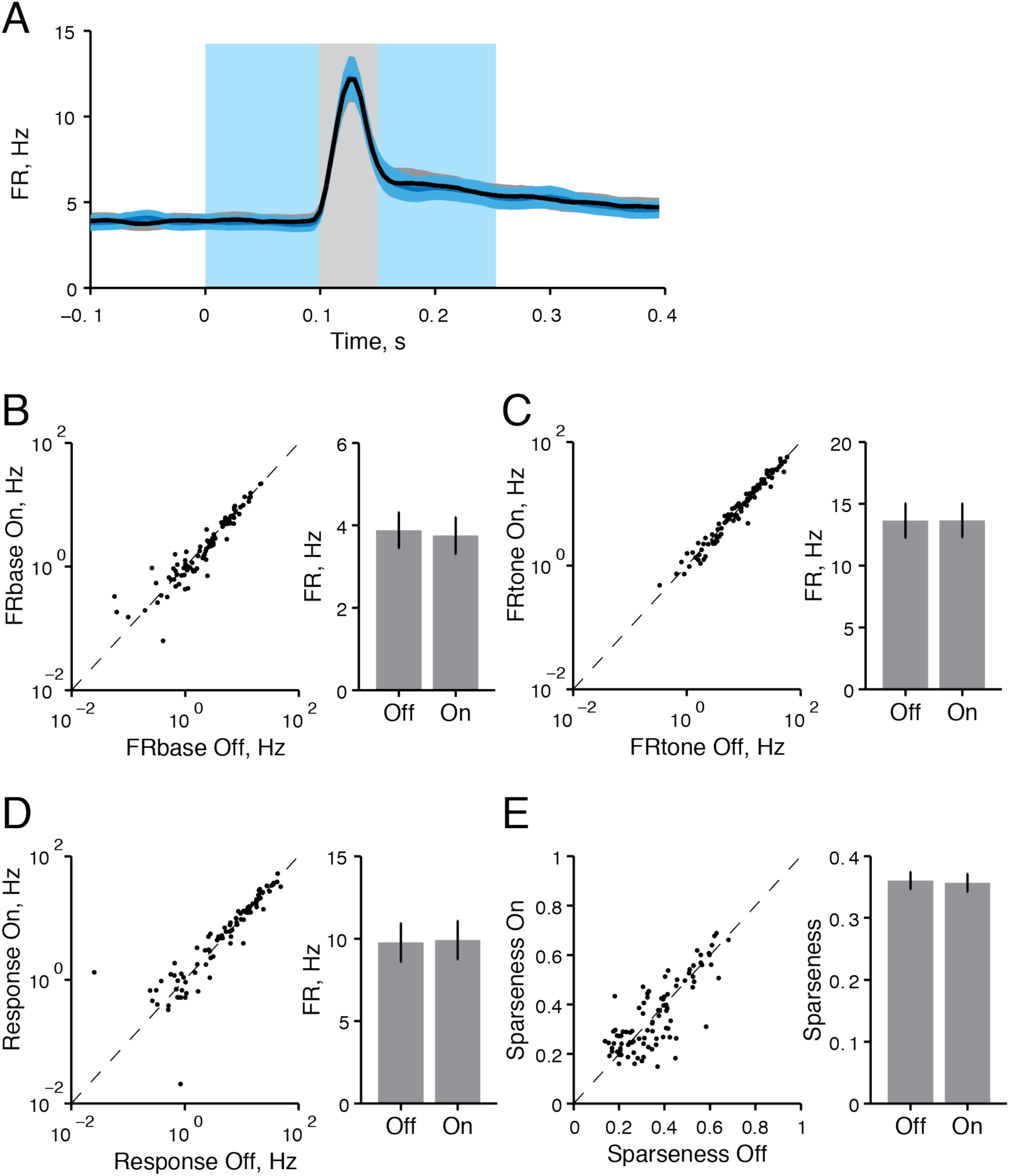
Photo-stimulation of BLA-TRN projections in mice expressing control virus does not affect spontaneous and tone-evoked activity in AC. **A.** Mean PSTH of the AC neurons in response to a tone on light-On (blue) and light-Off (black) trials. Plot shows data from all recorded neurons from 3 mice. Mean ± SEM. **B-E**. Photo-stimulation of BLA-TRN projections did not have a significant effect on spontaneous firing rate (FRbase, B, paired ttest, t_94_=1.58, p=0.12), tone-evoked activity (FRtone, C, paired ttest, t_94_=0.04, p=0.96), the amplitude of tone-evoked response (D, paired ttest, t_94_=0.46, p=0.65), or tuning sparseness of neurons in AC (E, paired ttest, t_94_=0.39, p=0.70). Left panel: Scatter plot of firing rate (B-D) or sparseness (E) on light-On plotted versus light-Off trials. Each circle represents a single unit. Right panel: Mean ± SEM of measures from the left panel.

**Figure S3.**
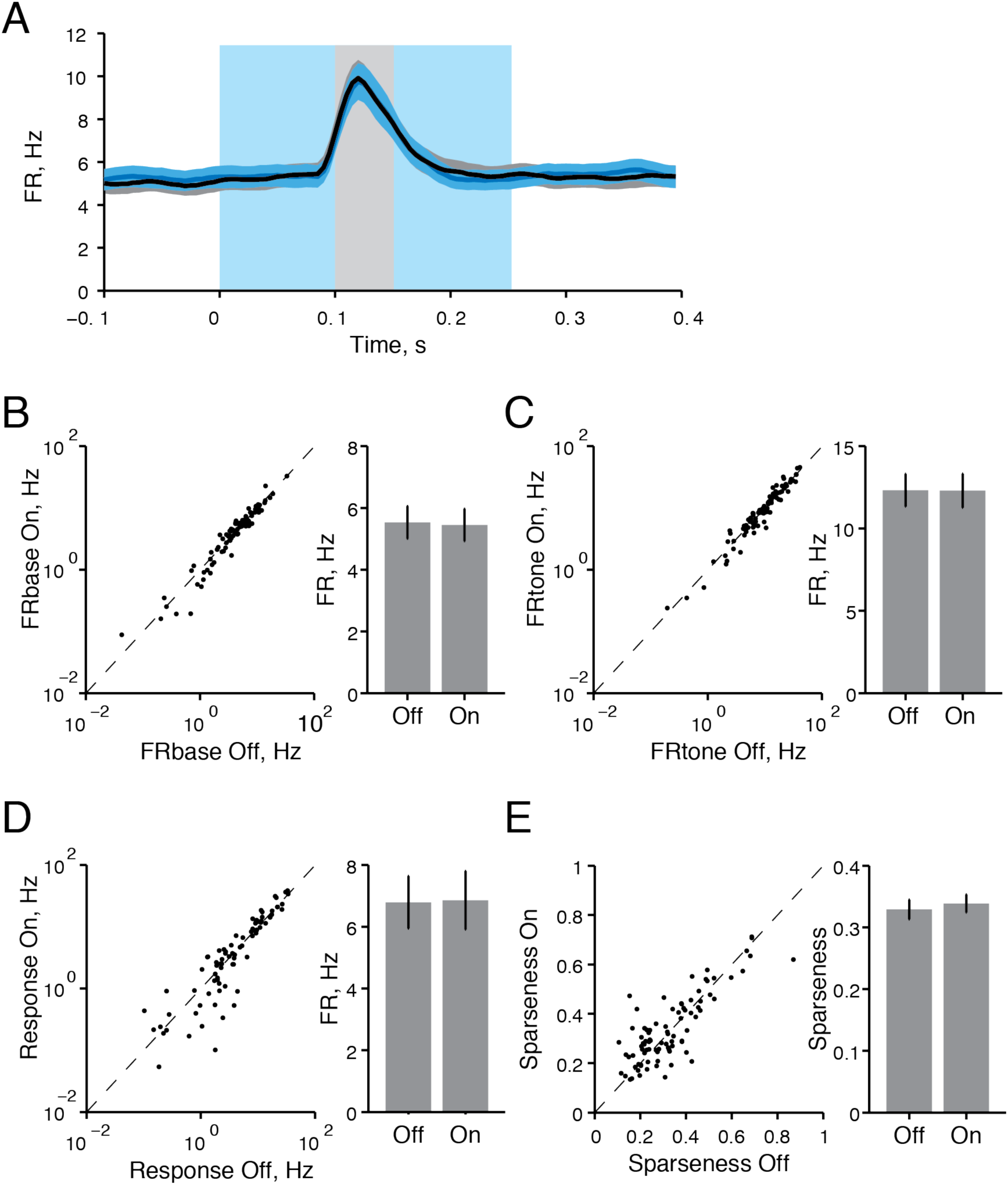
Photo-stimulation of BLA-TRN projections in mice expressing control virus does not affect spontaneous and tone-evoked activity in MGB. **A.** Mean PSTH of the MGB neurons in response to a tone on light-On (blue) and light-Off (black) trials. Plot shows data from all recorded neurons from 3 mice. Mean ± SEM. B-E. Photo-stimulation of BLA-TRN projections did not have a significant effect on spontaneous firing rate (B, FRbase, paired ttest, t_88_=0.59, p=0.56), tone-evoked activity (C, FRtone, paired ttest, t_88_=0.66, p=0.95), the amplitude of tone-evoked response (D, paired ttest, t_88_=0.19, p=0.85), or tuning sparseness of neurons in MGB (E, paired ttest, t_88_=1.03, p=0.31). Left panel: Scatter plot of firing rate (B-D) or sparseness (E) on light-On plotted versus light-Off trials. Each circle represents a single unit. Right panel: Mean ± SEM of measures from the left panel.

## References

1. Öhman, A., Flykt, A., and Esteves, F. (2001) Emotion drives attention: Detecting the snake in the grass. Journal of Experimental Psychology: General, 130(3): 466–478.

2. Phelps, E.A., Ling, S., and Carrasco, M. (2006) Emotion facilitates perception and potentiates the perceptual benefits of attention. Psychological science, 17(4): 292–299.

3. Aizenberg, M., Mwilambwe-Tshilobo, L., Briguglio, J.J., Natan, R.G., and Geffen, M.N. (2015) Bidirectional Regulation of Innate and Learned Behaviors That Rely on Frequency Discrimination by Cortical Inhibitory Neurons. PLoS biology, 13(12).

4. Li, W., Howard, J.D., Parrish, T.B., and Gottfried, J.A. (2008) Aversive learning enhances perceptual and cortical discrimination of indiscriminable odor cues. Science (New York, N.Y.), 319(5871): 1842–1845.

5. Resnik, J. and Paz, R. (2015) Fear generalization in the primate amygdala. Nature neuroscience, 18(2): 188–190.

6. Resnik, J., Sobel, N., and Paz, R. (2011) Auditory aversive learning increases discrimination thresholds. Nature neuroscience, 14(6): 791–796.

7. Ferrarelli, F. and Tononi, G. (2011) The thalamic reticular nucleus and schizophrenia. Schizophrenia bulletin, 37(2): 306–315.

8. Young, A. and Wimmer, R.D. (2017) Implications for the thalamic reticular nucleus in impaired attention and sleep in schizophrenia. Schizophrenia research, 180: 44–47.

9. Phillips, M.L., Drevets, W.C., Rauch, S.L., and Lane, R. (2003) Neurobiology of emotion perception II: Implications for major psychiatric disorders. Biological psychiatry, 54(5): 515–528.

10. LeDoux, J.E. (2000) Emotion circuits in the brain. Annu Rev Neurosci, 23: 155–84.

11. Quirk, G.J., Armony, J.L., and LeDoux, J.E. (1997) Fear conditioning enhances different temporal components of tone-evoked spike trains in auditory cortex and lateral amygdala. Neuron, 19(3): 613–24.

12. Ghosh, S. and Chattarji, S. (2015) Neuronal encoding of the switch from specific to generalized fear. Nature neuroscience, 18(1): 112–120.

13. Grewe, B.F., et al. (2017) Neural ensemble dynamics underlying a long-term associative memory. Nature, 543(7647): 670–675.

14. Grosso, A., Cambiaghi, M., Concina, G., Sacco, T., and Sacchetti, B. (2015) Auditory cortex involvement in emotional learning and memory. Neuroscience, 299: 45–55.

15. Kumar, S., von Kriegstein, K., Friston, K., and Griffiths, T.D. (2012) Features versus feelings: dissociable representations of the acoustic features and valence of aversive sounds. The Journal of neuroscience: the official journal of the Society for Neuroscience, 32(41): 14184–14192.

16. Sacco, T. and Sacchetti, B. (2010) Role of secondary sensory cortices in emotional memory storage and retrieval in rats. Science (New York, N.Y.), 329(5992): 649–656.

17. Weinberger, N.M. (2004) Specific long-term memory traces in primary auditory cortex. Nature reviews. Neuroscience, 5(4): 279–290.

18. Padmala, S. and Pessoa, L. (2008) Affective learning enhances visual detection and responses in primary visual cortex. J Neurosci, 28(24): 6202–6210.

19. Chavez, C.M., McGaugh, J.L., and Weinberger, N.M. (2009) The basolateral amygdala modulates specific sensory memory representations in the cerebral cortex. Neurobiology of learning and memory, 91(4): 382–392.

20. Letzkus, J.J., Wolff, S.B., Meyer, E.M.M., Tovote, P., Courtin, J., Herry, C., and Lüthi, A. (2011) A disinhibitory microcircuit for associative fear learning in the auditory cortex. Nature, 480(7377): 331–335.

21. Yang, Y., Liu, D.-Q.Q., Huang, W., Deng, J., Sun, Y., Zuo, Y., and Poo, M.-M.M. (2016) Selective synaptic remodeling of amygdalocortical connections associated with fear memory. Nature neuroscience, 19(10): 1348–1355.

22. Zikopoulos, B. and Barbas, H. (2012) Pathways for emotions and attention converge on the thalamic reticular nucleus in primates. The Journal of neuroscience: the official journal of the Society for Neuroscience, 32(15): 5338–5350.

23. Steriade, M., Deschênes, M., Domich, L., and Mulle, C. (1985) Abolition of spindle oscillations in thalamic neurons disconnected from nucleus reticularis thalami. Journal of neurophysiology, 54(6): 1473–1497.

24. Pinault, D. (2004) The thalamic reticular nucleus: structure, function and concept. Brain Research Reviews, 46(1): 1–31.

25. Ahrens, S., et al. (2015) ErbB4 regulation of a thalamic reticular nucleus circuit for sensory selection. Nature neuroscience, 18(1): 104–111.

26. Halassa, M.M., Chen, Z., Wimmer, R.D., Brunetti, P.M., Zhao, S., Zikopoulos, B., Wang, F., Brown, E.N., and Wilson, M.A. (2014) State-dependent architecture of thalamic reticular subnetworks. Cell, 158(4): 808–821.

27. Wimmer, R.D., Schmitt, L.I., Davidson, T.J., Nakajima, M., Deisseroth, K., and Halassa, M.M. (2015) Thalamic control of sensory selection in divided attention. Nature, 526(7575): 705–709.

28. Lerner, T.N., Ye, L., and Deisseroth, K. (2016) Communication in Neural Circuits: Tools, Opportunities, and Challenges. Cell, 164(6): 1136–1150. PMC5725393

29. Jhang, J., Lee, H., Kang, M.S., Lee, H.S., Park, H., and Han, J.H. (2018) Anterior cingulate cortex and its input to the basolateral amygdala control innate fear response. Nature communications, 9(1): 2744. PMC6048069

30. Znamenskiy, P. and Zador, A.M. (2013) Corticostriatal neurons in auditory cortex drive decisions during auditory discrimination. Nature, 497(7450): 482–5. 3670751

31. Shosaku, A. (1986) Cross-correlation analysis of a recurrent inhibitory circuit in the rat thalamus. Journal of neurophysiology, 55(5): 1030–1043.

32. Huguenard, J.R. (1996) Low-threshold calcium currents in central nervous system neurons. Annual review of physiology, 58: 329–48.

33. Willis, A.M., Slater, B.J., Gribkova, E.D., and Llano, D.A. (2015) Open-loop organization of thalamic reticular nucleus and dorsal thalamus: a computational model. J Neurophysiol, 114(4): 2353–67. PMC4620136

34. Gribkova, E.D., Ibrahim, B.A., and Llano, D.A. (2018) A novel mutual information estimator to measure spike train correlations in a model thalamocortical network. J Neurophysiol, 120(6): 2730–2744. PMC6337027

35. Traub, R.D., et al. (2005) Single-column thalamocortical network model exhibiting gamma oscillations, sleep spindles, and epileptogenic bursts. J Neurophysiol, 93(4): 2194–232.

36. Haas, J.S. and Landisman, C.E. (2011) State-dependent modulation of gap junction signaling by the persistent sodium current. Front Cell Neurosci, 5: 31. PMC3263475

37. Romanski and Ledoux. (1993) Somatosensory and Auditory Convergence in the Lateral Nucleus of the Amygdala. Somatosensory and Auditory Convergence in the Lateral Nucleus of the Amygdala.

38. Romanski, L.M. and LeDoux, J.E. (1992) Equipotentiality of thalamo-amygdala and thalamo-cortico-amygdala circuits in auditory fear conditioning. The Journal of neuroscience: the official journal of the Society for Neuroscience, 12(11): 4501–4509.

39. Campeau, S. and Davis, M. (1995) Involvement of the central nucleus and basolateral complex of the amygdala in fear conditioning measured with fear-potentiated startle in rats trained concurrently with auditory and visual conditioned stimuli. The Journal of neuroscience: the official journal of the Society for Neuroscience, 15(3 Pt 2): 2301–2311.

40. Goosens, K.A. and Maren, S. (2001) Contextual and auditory fear conditioning are mediated by the lateral, basal, and central amygdaloid nuclei in rats. Learning & memory (Cold Spring Harbor, N.Y.), 8(3): 148–155.

41. Antunes, R. and Moita, M.A. (2010) Discriminative auditory fear learning requires both tuned and nontuned auditory pathways to the amygdala. The Journal of neuroscience: the official journal of the Society for Neuroscience, 30(29): 9782–9787.

42. Boatman, J.A. and Kim, J.J. (2006) A thalamo-cortico-amygdala pathway mediates auditory fear conditioning in the intact brain. The European journal of neuroscience, 24(3): 894–900.

43. Johansen, J.P., Hamanaka, H., Monfils, M.H., Behnia, R., Deisseroth, K., Blair, H.T., and LeDoux, J.E. (2010) Optical activation of lateral amygdala pyramidal cells instructs associative fear learning. Proceedings of the National Academy of Sciences of the United States of America, 107(28): 12692–12697.

44. Aizenberg, M. and Geffen, M.N. (2013) Bidirectional effects of aversive learning on perceptual acuity are mediated by the sensory cortex. Nature neuroscience, 16(8): 994–996.

45. Kimura, A., Imbe, H., Donishi, T., and Tamai, Y. (2007) Axonal projections of single auditory neurons in the thalamic reticular nucleus: implications for tonotopy-related gating function and cross-modal modulation. European Journal of Neuroscience, 26(12): 3524–3535.

46. Pham, T. and Haas, J.S. (2018) Electrical synapses between inhibitory neurons shape the responses of principal neurons to transient inputs in the thalamus: a modeling study. Scientific reports, 8(1): 7763. PMC5958104

47. Guillery, R.W., Feig, S.L., and Lozsadi, D.A. (1998) Paying attention to the thalamic reticular nucleus. Trends in neurosciences, 21(1): 28–32.

48. Jones, E.G. (1975) Some aspects of the organization of the thalamic reticular complex. Journal of Comparative Neurology.

49. Kimura, A., Yokoi, I., Imbe, H., Donishi, T., and Kaneoke, Y. (2012) Auditory thalamic reticular nucleus of the rat: anatomical nodes for modulation of auditory and cross-modal sensory processing in the loop connectivity between the cortex and thalamus. The Journal of comparative neurology, 520(7): 1457–1480.

50. Pinault, D. and Deschênes, M. (1998) Projection and innervation patterns of individual thalamic reticular axons in the thalamus of the adult rat: A three-dimensional, graphic, and morphometric analysis. Journal of Comparative Neurology.

51. Villa, A.E.P. (1990) Physiological differentiation within the auditory part of the thalamic reticular nucleus of the cat. Brain Research Reviews.

52. Crick, F. (1984) Function of the thalamic reticular complex: the searchlight hypothesis. Proceedings of the National Academy of Sciences, 81(14): 4586–4590.

53. Landisman, C.E., Long, M.A., Beierlein, M., Deans, M.R., Paul, D.L., and Connors, B.W. (2002) Electrical synapses in the thalamic reticular nucleus. J Neurosci, 22(3): 1002–9.

54. Brown, J.W., Taheri, A., Kenyon, R.V., Berger-Wolf, T., and Llano, D.A. (2019) A computational model of intrathalamic signaling via open-loop thalamo-reticular-thalamic architectures. BioRXiv, 10.1101/574178.

55. Lennartz and Weinberger. (1992) Frequency-Specific Receptive Field Plasticity in the Medial Geniculate Body Induced by Pavlovian Fear Conditioning Is Expressed in the Anesthetized Brain. Behavioral Neuroscience.

56. Weinberger, N.M., Javid, R., and Lepan, B. (1993) Long-term retention of learning-induced receptive-field plasticity in the auditory cortex. Proceedings of the National Academy of Sciences of the United States of America, 90(6): 2394–2398.

57. Bakin, J.S. and Weinberger, N.M. (1990) Classical conditioning induces CS-specific receptive field plasticity in the auditory cortex of the guinea pig. Brain Res, 536(1-2): 271–86.

58. Moczulska, K.E., Tinter-Thiede, J., Peter, M., Ushakova, L., Wernle, T., Bathellier, B., and Rumpel, S. (2013) Dynamics of dendritic spines in the mouse auditory cortex during memory formation and memory recall. Proceedings of the National Academy of Sciences of the United States of America, 110(45): 18315–18320.

59. Amaral, D.G. and Price, J.L. (1984) Amygdalo cortical projections in the monkey (Macaca fascicularis). Journal of Comparative Neurology, 230(4): 465–496.

60. Yukie, M. (2002) Connections between the amygdala and auditory cortical areas in the macaque monkey. Neuroscience research, 42(3): 219–229.

61. Wood, K.C., Blackwell, J.M., and Geffen, M.N. (2017) Cortical inhibitory interneurons control sensory processing.. Current Opinion in Neurbiology.

